# Influence of past climate changes and human-mediated introduction on patterns of genetic diversity distribution of the Papuan nutmeg (*Myristica argentea*) Warb

**DOI:** 10.1101/2025.02.05.636754

**Authors:** Jakty Kusuma, Adinda C. Milaba, Amalia Paramitha, Victoria Malaval, Mei Nita Sari, Cahya Nurhuda, Marie Couderc, Nora Scarcelli, John Watts, Jérôme Duminil

## Abstract

Understanding the distribution of genetic diversity of endemic food tree species holds significant importance for conservation planning and the formulation of effective management strategies. Natural and anthropogenic factors can influence species’ geographic range and distribution of genetic diversity. Here, we tested the relative influence of barriers to gene flow (rivers, mountains and past climate change) and recent human activities on the evolutionary history of an endemic species from Fakfak district, Papua Island, *Myristica argentea* Warb (Myristicaceae), better known as Papuan nutmeg. *Myristica argentea* is a spice tree species used by local populations since centuries/millennia. The species has been introduced in other locations from the region. Fourteen microsatellite markers (nSSRs), whole chloroplast (cp) genome, and nuclear-targeted sequence genes (nrDNA sequence) were used to characterize the genetic diversity, population structure, and demographic history of the species across its distribution range (native and introduced populations). We found that native populations of the species in Fakfak present higher levels of genetic diversity than introduced populations. We further detected the presence of a moderate genetic structure (*F*_ST_ = 0.143), and the presence of three intraspecific genetic clusters. Additionally, our findings indicate that population fragmentation within the native range of the species in Fakfak, may have been influenced by the presence of riparian networks, mountain ranges and by Pleistocene climatic fluctuations. Our results suggest that all introduced populations were sourced mainly from the native population of Raduria, in the South of Fakfak region. Given the population genetic variation across the ranges of Papuan nutmeg, management plans should not treat them as single populations, but rather consider the broader genetic diversity within its range.

## INTRODUCTION

Islands offer a distinct narrative of species’ genetic variation and diversification when compared to their continental counterparts. Despite numerous studies that have assessed evolutionary history at a continental level (Bell et al., 2018; Couvreur et al., 2021; Graham, 2018; Helmstetter et al., 2020; Lu et al., 2018), our understanding of evolutionary dynamics at the island level remains scarce. This knowledge gap is remarkable given the central role that islands play in biodiversity and their enormous contribution to global diversity (Condamine et al., 2017).

The Indonesian archipelago is part of the Malesian biogeographic region and consists of ∼17,000 islands (Raes & Van Welzen, 2009; von Rintelen et al., 2017). It is a highly biodiverse region with various climate and unique dynamic geotectonic activities (Joyce et al., 2020), which are drivers of plant diversity. Papua Island (internationally known as New Guinea Island), situated in the eastern part of Indonesia, is renowned for being home to one of the most diverse and abundant flora in the world (Cámara-Leret et al., 2020). An estimated 90% of the tree species found on Papua Island are unique and endemic to the region, showcasing its exceptional biodiversity and ecological significance (Barstow et al., 2022). Thus, Papua Island is home to species-rich genera that contain more than 80 species per genus (Cámara-Leret et al., 2020)

The high number of tree species on Papua Island has been attributed to its highly variable topography, a complex geological history and steep environmental gradients (Morley, 2000; Slik et al., 2015), as well as its historical prints of glaciation (Wallace, 1878). It has been argued that during the glacial event (Last Glacial Maximum), many forest-restricted species persisted in localized regions, referred to as forest refugia, where relatively favorable climatic conditions allowed for their survival (Blatrix et al., 2013; Hewitt, 1996; Pouteau et al., 2015). These forest refugia acted as reservoirs of genetic diversity and, as the climate warmed during the Holocene, these populations expanded and recolonized other areas (Hewitt, 2000). As a result, the populations that survived in these refugia might have experienced a drastic change in population size as well as a rupture of gene flow among refugia. This is expected to lead to population differentiation, as geographic barriers are commonly recognized as a key mechanism driving population divergence. Mountains, along with rivers, are one of the main topographic factors associated with long-term barriers to gene flow (Khanal et al., 2018; Sánchez-Montes et al., 2018). As the Papua Island topography is highly complex (Baldwin et al., 2012; Toussaint et al., 2021), it is thus interesting to test the forest refugia hypothesis on this island, which remains poorly documented.

Papua Island is home to the highest number of *Myristica* species (de Wilde, 2014), which suggest that this area is the center of origin and diversification of the genus. *Myristica argentea* Warb.—known as Papuan nutmeg, Fakfak nutmeg, or Macassar nutmeg—is one of the most economically and ethnobotanically recognized species from Papua Island. The species is natively distributed in West Papua, Fakfak peninsula (de Wilde, 2014). It is commonly used as a source of nutmeg and mace in the same ways as the Banda nutmeg (*Myristica fragrans*).

There is evidence that the natural distribution of *M. argentea* is confined to a limited geographic range (de Wilde, 1995, 2014), and it is commonly observed that species with restricted distributions tend to display a low levels of genetic diversity (DeJoode & Wendel, 1992; Frankham, 1996, 1997). The restricted genetic diversity observed in narrowly distributed island taxa, as demonstrated by previous meta-analyses (González et al., 2020), is thought to be influenced by various factors that remain a subject of debate (Frankham, 1997). Some proposed explanations include the effect of limited dispersal abilities (Hindley et al., 2018) and small population sizes (Barret, 1996; Frankham, 1996). However, several examples explained that insular species with endemic status exhibit higher genetic diversity compared with their continental lineages (García-Verdugo et al., 2015; Salmona et al., 2023).

Furthermore, it is also noteworthy that anthropogenic activities can influence the distribution range and the level of genetic diversity of species. For instance, *M. argentea* has been introduced to neighboring islands outside its native range due to its similarity to *M. fragrans*. This led to the expansion of Papuan nutmeg’s distribution range from the Fakfak district to nearby islands such as Ambon and Seram in the Moluccas archipelago. Therefore, it is interesting to investigate the levels of genetic diversity within both the restricted native range and the introduced populations of this species.

Against the above background, our objectives here were: (i) to investigate the influence of topographic (mountains, rivers) and past climate changes on the patterns of evolutionary history of Papuan nutmeg by characterizing the distribution of genetic diversity in its native range; (ii) to investigate the influence of human-mediated dispersal on the genetic structure of the species sby characterizing the distribution of genetic diversity of introduced populations of Papuan nutmeg.

## MATERIALS AND METHODS

### Study area, plant materials and geographic sampling

*Myristica argentea* is native from Fakfak region, Papua Island and primarily grows in forests below 250 meters above sea level (Musaad et al., 2016; Wahyuni & Bermawie, 2020; de Wilde, 2014). The island of Papua itself is a relatively young landmass that has emerged and developed over the last 25 - 2 million years (Evans, 2020), during which it was colonized by numerous species (Oliver et al., 2017; Toussaint et al., 2014). We collected in the field 468 individuals of *M. argentea* (Figure 1 and Supplementary file S1). These samples originated from fifteen populations in Papua Island (thirteen from Fakfak Peninsula, and one from Manokwari and Raja Ampat) and two populations from South Moluccas archipelago (Ambon and Seram Island). The thirteen populations from Fakfak peninsula are located in the region of origin of the species (called ‘native’ populations hereafter), whereas additional populations (Manokwari, Raja Ampat, Seram and Ambon) correspond to human-driven introductions outside of the original distribution of the species (called ‘introduced’ populations hereafter). Interviews with farmers informed us that native populations are composed of a mix of wild individuals, that were kept during land cleaning (the oldest individuals of the population), and planted individuals, originated from seeds of wild individuals from the same location.

**Figure 1.**
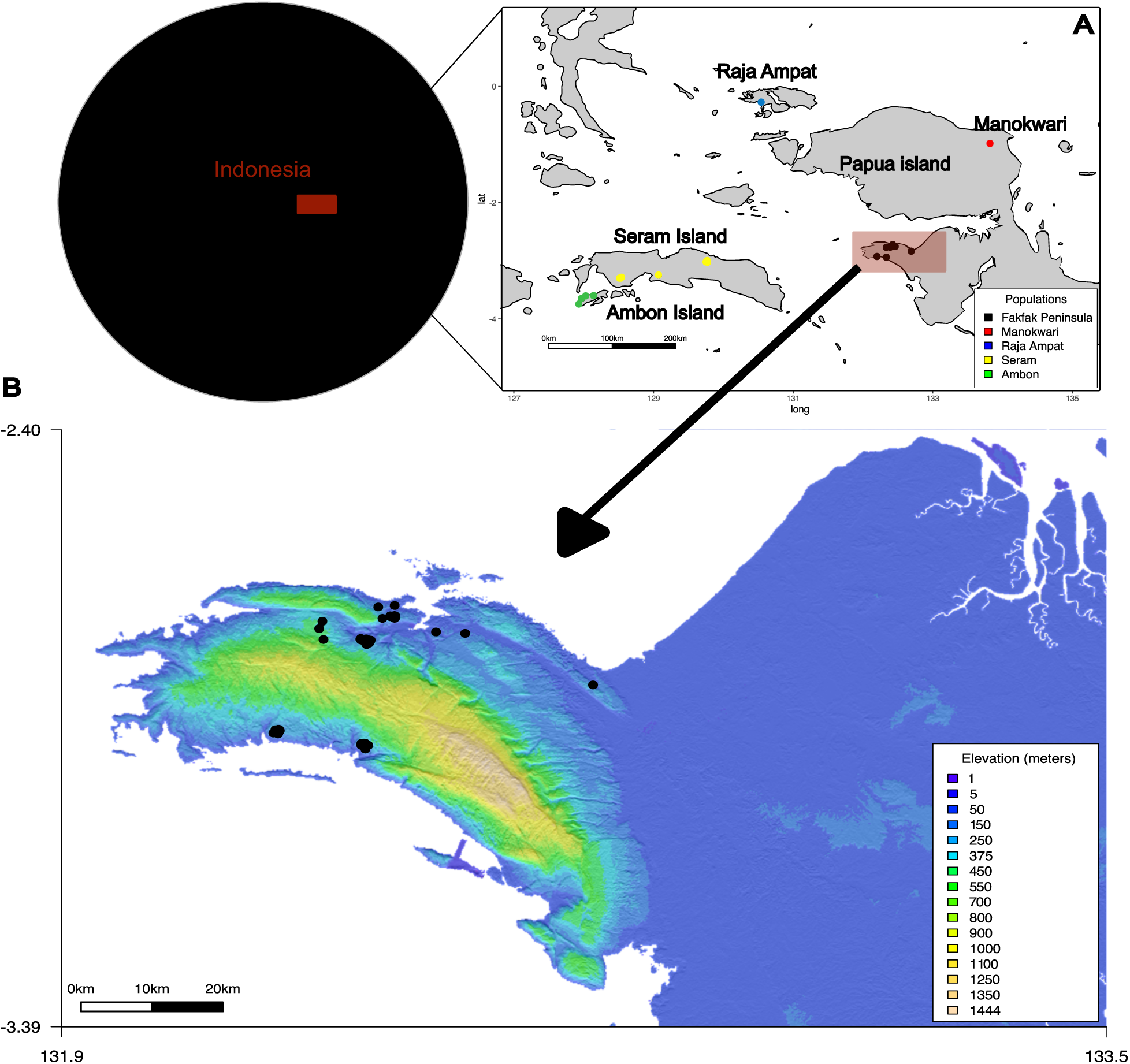
The inner caption within Indonesian archipelago is pointing to (A) sampling locations of *Myristica argentea* populations in Indonesia; (B) sampling locations of *M. argentea* subpopulations in Fakfak Peninsula population. Individuals are indicated by a color according to their population.

### Molecular genetic analysis

*DNA extraction and nuclear microsatellite markers.* For each sample, young fresh leaves were collected and immediately dried in silica gel. DNA was extracted from dried leaves using the protocol of Mariac *et al*. (2006). Samples were genotyped using 14 nuclear microsatellite markers (nSSRs) following PCR conditions as described in Kusuma *et al*. (2020). Two of the primer pairs (*Myr*28 and *Myr*42) amplified two different genomic regions, so we split them into four different markers (namely *Myr*28a and *Myr*28b, *Myr*42a and *Myr*42b). Genotyping was done on an ABI 3500 XL and analyzed using Geneious Prime® 2023.0.4 (2012a), with manual corrections.

*Chloroplast DNA marker.* A subsample of 43 *M. argentea* individuals (3 to 5 individuals per population) and three individuals of the sister species *M. fragrans*, used as outgroups, were sequenced (*N*_TOTAL_ = 46; Supplementary File S1). Whole genome library constructions were done following Mariac *et al* (2014). Then raw data were analyzed following the method explained in Scarcelli et al. (2006). Briefly, we cleaned and filtered the reads using DEMULTADAPT (https://github.com/Maillol/demultadapt) and CUTADAPT 1.2.1 (Martin, 2011), then we mapped them with BWA (Li & Durbin, 2009) using the *M. argentea* cpDNA genome as reference (Kusuma et al., 2023; GenBank accession: OP866724.1). We called SNPs using VARSCAN ver. 2.3.7 (Koboldt et al., 2012) and filtered them using VCFtools (Danecek et al., 2011). All scripts used are described in Supplementary File S2.

*Nuclear targeted-sequence marker.* We used the same individuals as for cpDNA analysis (*N* = 46, see Supplementary File S1). Genomic libraries were prepared in ARCAD facilities (Montpellier, France). We then used targeted-capture strategies using a baiting kit of 469 nuclear genes, initially developed for a sister family of Myristicaceae, the Annonaceae (Couvreur et al., 2019). We then processed the DNA sequences of the 46 individuals with the pipeline SECAPR v.1.14 (Andermann et al., 2018). This method is beneficial for identifying SNPs because it generates a pseudo-reference made from the target regions’ sequences, except for the paralog regions, which were identified and taken out automatically. BWA v.0.7.12 (Li & Durbin, 2009) was used for mapping and GATK v.4 (McKenna et al., 2010) was used to remove duplicates and to call SNPs. We filtered the data by keeping only biallelic SNPs with a mapping quality >40%, depth >25, and quality by depth >2 using BCFtools (Li, 2011). Additionally, we removed SNPs with allele frequency <0.01 and excluded monomorphic sites. Scripts used for this analysis are available in https://github.com/jaktykusuma/myristica/tree/master/argentea.

### Genetic diversity and population structure

Being highly polymorphic, nSSRs are markers of choice to assess the distribution of the genetic diversity within populations. We characterized the levels of genetic diversity within population (*N*_POP_ = 17) using SPAGeDI 1.5 (Hardy & Vekemans, 2002) with the following indices: number of alleles (*N*_A_), observed and expected heterozygosity (*H*_O_ and *H*_E_, respectively), and the inbreeding coefficient (*F*_IS_). We calculated the rarefied allelic richness (*A*_R_) using R (R Core Team, 2022) package *‘hierfstat’* (Goudet, 2005). Presence of null alleles (*F*_null_) and the *F*_IS_ corrected for the presence of null alleles were calculated using INEST 2.2 (Chybicki & Burczyk, 2009). Levels of genetic diversity was also estimated for each population using the cpDNA dataset. Thus, we determined the number of haplotypes (*H*), nucleotide diversity (π), number of segregating sites (S), and number of parsimony-informative sites per population with DnaSP 6.12.03 (Librado & Rozas, 2009).

Using SSR, we characterized the population structure by getting estimates of pairwise *F*_ST_ between populations using SPAGeDI 1.5 (Hardy & Vekemans, 2002), and by applying a Bayesian clustering approach as implemented in STRUCTURE (Pritchard et al., 2000). In addition, we conducted a population structure analysis using the nrDNA sequence dataset using fastStructure, and used the resulting genetic groups in subsequent demographic analyses. We also visualized the relationships and connections between cpDNA haplotypes within and among populations by reconstructing an haplotype network using the maximum parsimonious tree method (Kannan & Wheeler, 2012) as implemented in PopART (Leigh & Bryant, 2015), with a 95% statistical parsimony criterion.

### Inference of population size

In order to test the presence of demographic changes in relation to past climate oscillations, we used the nrDNA sequences data to infer the evolution of native population size through time. The nrDNA sequences are adequate to conduct this analysis given their high potential to discriminate different evolutionary scenarios (Cornuet et al., 2010) and to improve the accuracy of parameters estimations (Smith & Flaxman, 2020). Using the nrDNA sequences, we utilized the Stairway Plot 2 approach (Liu & Fu, 2020), a model-flexible method that employs site-frequency spectra (SFS), to determine changes in effective population size (*N*_E_) over time within each genetic cluster. To ensure the accuracy of our results, we did not apply minor-allele frequency filter, as it could potentially bias the SFS. Since Stairway Plot relies on SNP counts for *N*_E_ inference, the exclusion of SNPs with missing data may introduce count biases. To mitigate this, we calculated the SFS for each intra-specific cluster observed within the native distribution of the species (as revealed by the Bayesian clustering analysis conducted on the nrDNA sequences dataset) following the methodology of Burgarella *et al*. (2018), i.e. by considering the minor-allele frequency at each SNP, multiplied by the mean number of haploid samples. This resulted in a new SFS that incorporated all observed site frequencies while minimizing SNP removal. We utilized two-thirds of the sites and performed 200 bootstraps for robustness. The number of observed sites was determined as the total length of the SECAPR pseudo-reference (see above) for each population, ensuring comprehensive coverage for our analysis. We applied a time calibration by using the Angiosperm wide mutation rate of 5.35e^−9^ mutations per site per year (De La Torre et al., 2017) and a generation time of 50 years.

### Population history

As an additional test of the influence of past-climate events, we inferred the ancestry and divergence time of the genetic clusters from nrDNA data, defined by STRUCTURE results, using DIYABC Random Forest 1.1.1 (Collin et al., 2021). The populations assigned to a given genetic cluster according to STRUCTURE results were pooled as a single genetic group to reduce demographic model complexity. We only kept individuals with a cluster assignment probability higher than 0.85.

We tested different scenarios (Supplementary Figure S1) of population divergence using various historical demographic parameters (Supplementary Table S1), without considering any admixture event. Coalescent simulations were performed for each scenario, with a total of 100,000 simulations. To assess the agreement between simulated and observed data, Principal Component Analysis (PCA) was conducted on the summary statistics from simulated data and the observed data. The study employed Approximate Bayesian Computation (ABC) with random forest (RF) in ‘*abcranger*’ module to perform scenario choice and parameter estimation (Pudlo et al., 2016; Raynal et al., 2019). The best scenarios were selected based on a classification vote, and RFs of 1,000 trees were used to estimate median point estimates and a 95% posterior credible interval for each demographic parameter.

## RESULTS

### Genetic diversity

In the nSSR dataset, we identified two loci (*Myr*12 and *Myr*26) that had more than 40% of missing data and 53 individuals had more than 30% of missing data at different populations (Supplementary Figure S2). The corresponding loci and individuals were removed, resulting in a dataset of 420 individuals on 14 nSSR diploid genotypes. The 14 nSSRs revealed a total of 137 alleles, with a mean number of alleles per population ranging from 2.57 (Raja Ampat) to 6.57 (Mamur) (see Table 1 for detailed values). Using allelic richness as an indicator of genetic diversity, we found that half of the populations within the species’ native range exhibited significantly higher levels when compared to the introduced population (Table 1). Specifically, a significantly higher allelic richness across loci was observed in Perwasak (5.58), Mamur (3.16), Pawada (3.40), Kokas (3.39), Nika (3.36), Ubadari (3.52), and Patimburak (3.31) compared to Ambon (3.09), Raja Ampat (2.08), Manokwari (2.43), and Seram (3.03) (ANOVA: P < 0.01). This finding suggests a greater genetic diversity in most of the native populations when compared to introduced populations. We found low levels of genetic diversity in Raja Ampat and Manokwari, as indicated by the rarefied allelic richness (*AR*_(*k* = 7)_; Table 1).

**Table 1.**
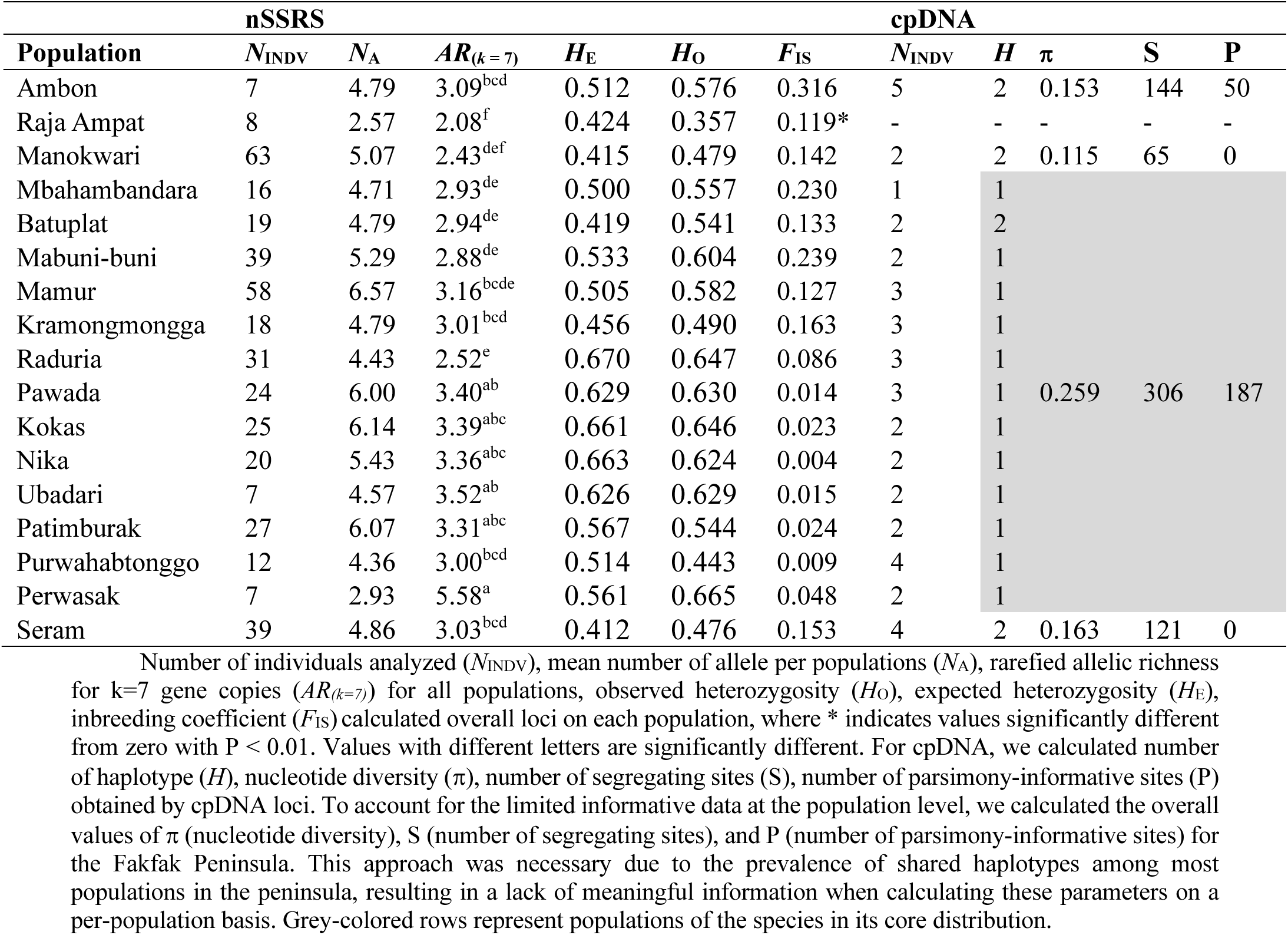
Genetic diversity of *Myristica argentea* inferred by 14 nSSRs in all populations.

### Population structure

Pairwise *F*_ST_ of nSSR dataset is between populations ranged from 0.008 (between Ambon and Seram) to 0.327 (between Raja Ampat and Perwasak; Supplementary Figure S3). Furthermore, the Bayesian clustering analysis for all populations revealed the presence of three distinct genetic clusters (*K* = 3; Figure 2). Based on individual assignment scores (> 0.85), we allocated each population to its corresponding genetic cluster. The first genetic cluster comprised Ambon, Raja Ampat, Manokwari, and Raduria populations. The second genetic cluster consisted of Batuplat and Mabuni-buni populations. The third genetic cluster included Mamur, Kramongmongga, Pawada, Kokas, Nika, Ubadari, and Patimburak populations. Mbahambandara, Purwahabtonggo, and Perwasak populations were considered as admixed populations. Based on these clustering, the native populations were assigned to three different geographically-structured genetic clusters. We found that introduced populations were grouped together with the native population of Raduria. We found a similar structuration pattern with the nrDNA sequence data (Supplementary Figure S4).

**Figure 2.**
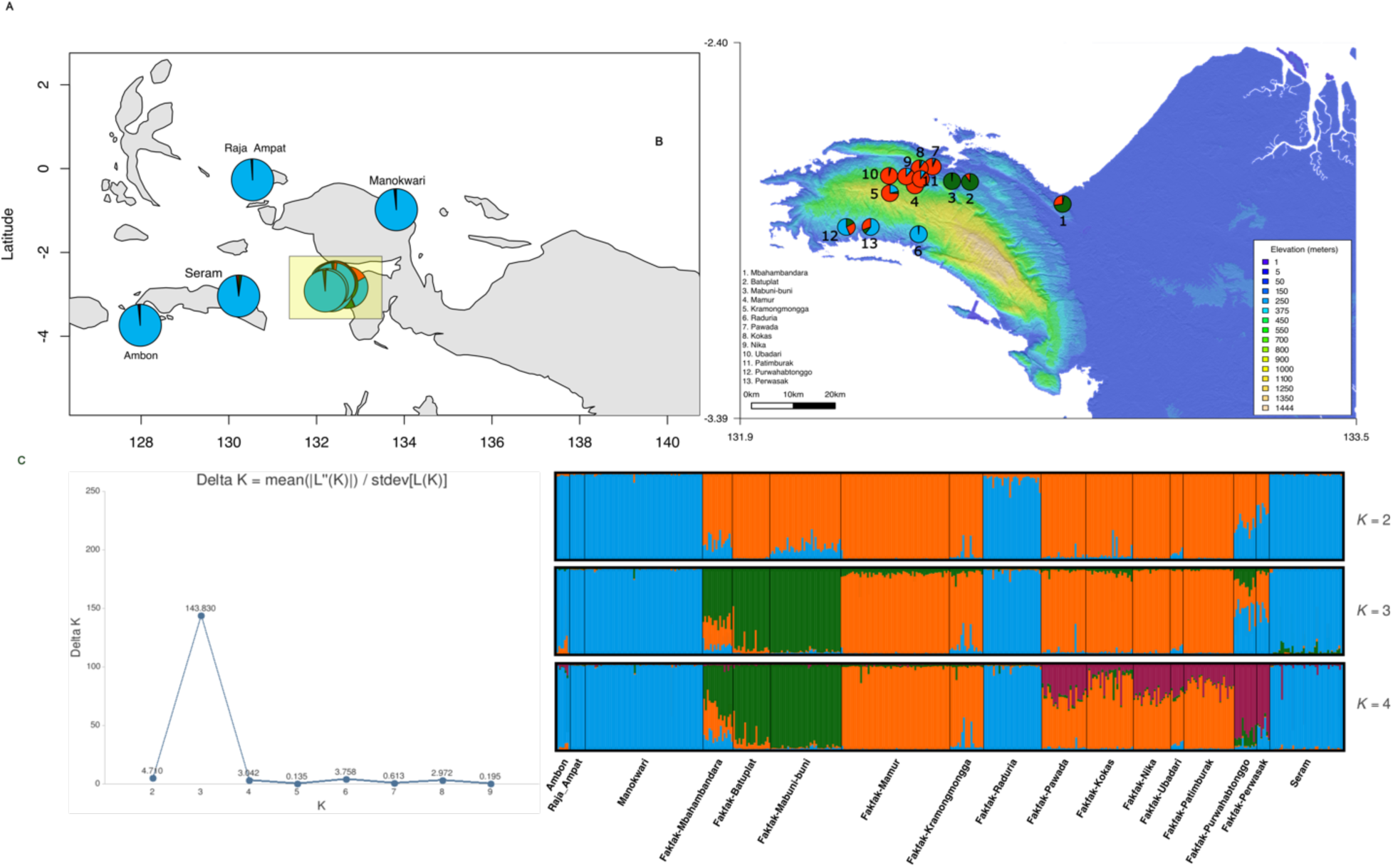
Population structure based on nSSRs in all seventeen populations using STRUCTURE. Geographical distribution of individuals for (A) *K* = 4 in all seventeen populations, and (B) in the species’ core distribution, Fakfak Peninsula. Delta *K* value (C), and barplot of individual admixture proportions for each individual inferred by STRUCTURE for *K* = 2 to *K* = 5 (D).

We detected 405 SNP, corresponding to twelve different cpDNA haplotypes within our *M. argentea* sampling (Figure 3). All native populations were fixed for one given haplotype, except for the Batuplat population, which displayed two distinct haplotypes (*Hap*4 and *Hap*12). Within native populations, one shared haplotype (*Hap*1) was present in five populations of Fakfak Peninsula (Figure 3A). The distribution of the haplotypes showed that the two haplotypes in Ambon (*Hap*4 and *Hap*7) and Seram (*Hap*7 and *Hap*10) were shared with the population from its native range, which includes Batuplat (*Hap*4), Raduria (*Hap*7), and Kokas (*Hap*10).

**Figure 3.**
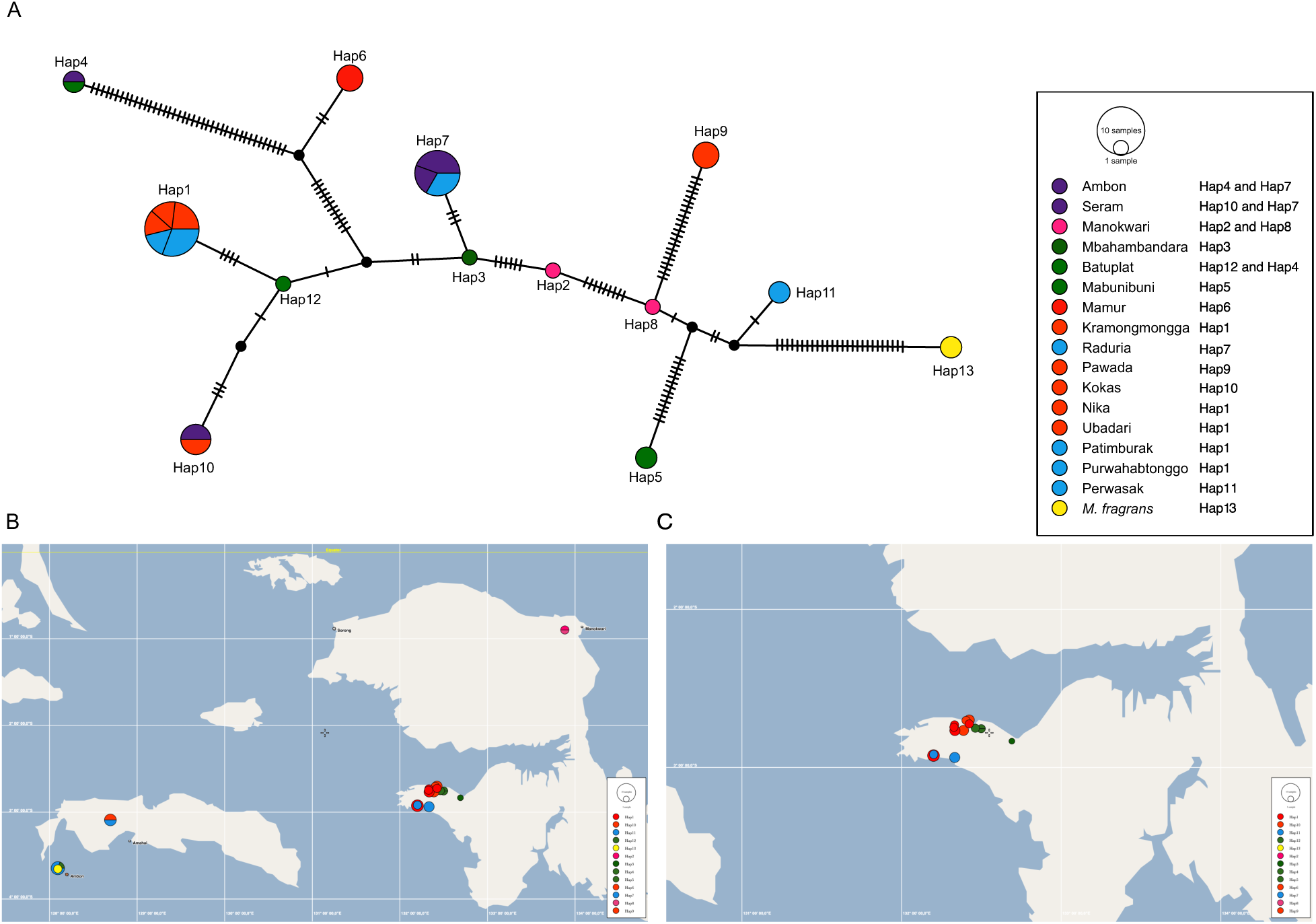
Maximum parsimonious network based on the cpDNA haplotypes. Each circle represents a unique haplotype according to their populations. The haplotypes were color-coded based on their geographical origin. Purple represents Ambon and Seram, pink represents Manokwari, and yellow represents the outgroup (*M. fragrans*). The color coding in Fakfak Peninsula corresponds to structure analysis. Cluster 1 (dark green) includes Mbahambandara, Mabuni-buni, and Batuplat. Cluster 2 (orange) consists of Mamur, Kramongmongga, Pawada, Kokas, Nika, Ubadari, and Patimburak. Cluster 3 (light blue) is composed of Raduria, Perwasak, and Purwahabtonggo. The legend provides the assignment of haplotypes in each population.

### Inference of population size

We observed three intra-specific clusters using the Bayesian clustering analysis conducted on the nrDNA sequences dataset (Figure 4): Cluster 1 included Mbahambandara, Mabuni-buni, and Batuplat populations (*N* = 5), Cluster 2 included Mamur, Kramongmongga, Pawada, Kokas, Nika, Ubadari, and Patimburak populations (*N* = 17) and Cluster 3 included Raduria, Perwasak and Purwahabtonggo populations (*N* = 9). The stairway plot analysis showed that the first two clusters underwent an expansion approximately 500,000 years ago (Figure 4A and 4B). Following this expansion, Cluster 1 exhibited a constant population size, as evidenced by the upper and lower bounds of the 95% confidence interval. Although the statistical support for this observation was not particularly strong (*P* > 0.005), the historical constant size persisted. In contrast, Cluster 2 experienced multiple pulses of population change, with a significant decline starting from approximately ∼20,000 to ∼7,500 years ago. However, after this decline, the population size remained constant. Regarding Cluster 3 (Figure 4C), a severe population decline was detected around ∼100,000 years ago, followed by a sudden expansion around ∼50,000 years ago. Subsequently, the population size stabilized and remained constant.

**Figure 4.**
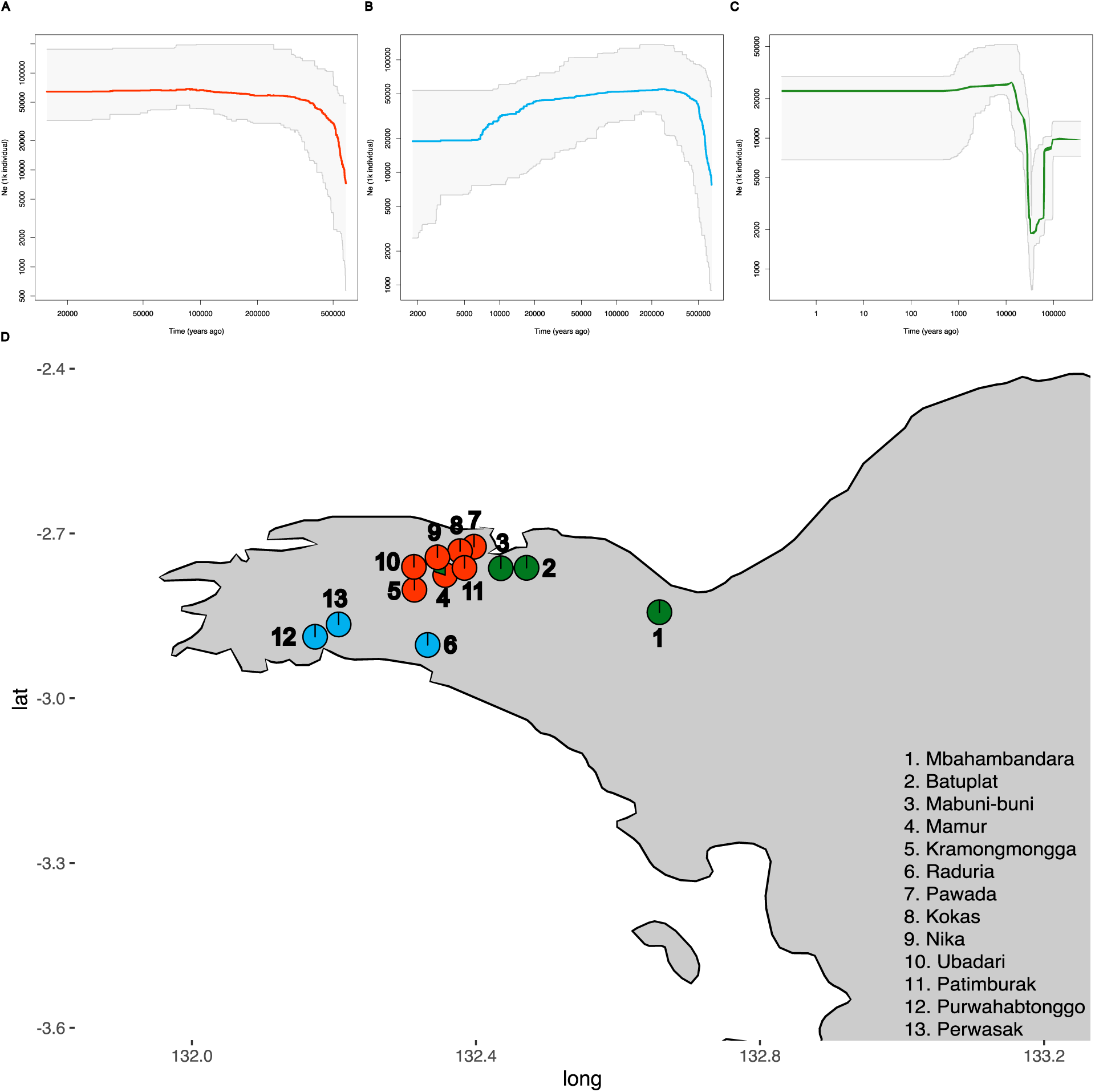
Inference of population histories produced by Stairway Plot on the nrDNA sequence in Fakfak subpopulations. (A) Cluster 1 (red dots in the map), including Mbahambandara, Mabuni-buni, and Batuplat; (B) Cluster 2 (blue dots), including Mamur, Kramongmongga, Pawada, Kokas, Nika, Ubadari, and Patimburak; and (C) Cluster 3 (green dots), including Raduria, Perwasak and Purwahabtonggo. Median values of *Ne* is indicated in solid colored line, while dashed lines correspond to the upper and lower bounds of the 95% confidence intervals. Original plots generated by Stairway Plot and confidence interval values in table format are available in Dryad (https://doi.org/10.5061/dryad.f7m0cfz1w). (D) Geographical distribution on each cluster.

### Population history

The PCA plots of both observed and simulated data were visually inspected (see Supplementary Figure S5). Our findings suggested that the considered models can accurately replicate the observed patterns of genetic diversity. Therefore, we deemed the proposed scenarios and prior combinations sufficient for conducting ABC analysis. Model selection according to DIYABC-RF favors scenario 1 as the best supported scenario (Figure 5), with a posterior probability of 0.783 and prior error of 0.35 and a Bayes factors of 2.889. This scenario suggested that all populations split at the same time from a single ancestral population. The divergence time was estimated at 201 to 417 generations ago, with median of 276 generations ago. Assuming 50 years of generation time, the split would fall around 10,500 to 20,850 years ago.

**Figure 5.**
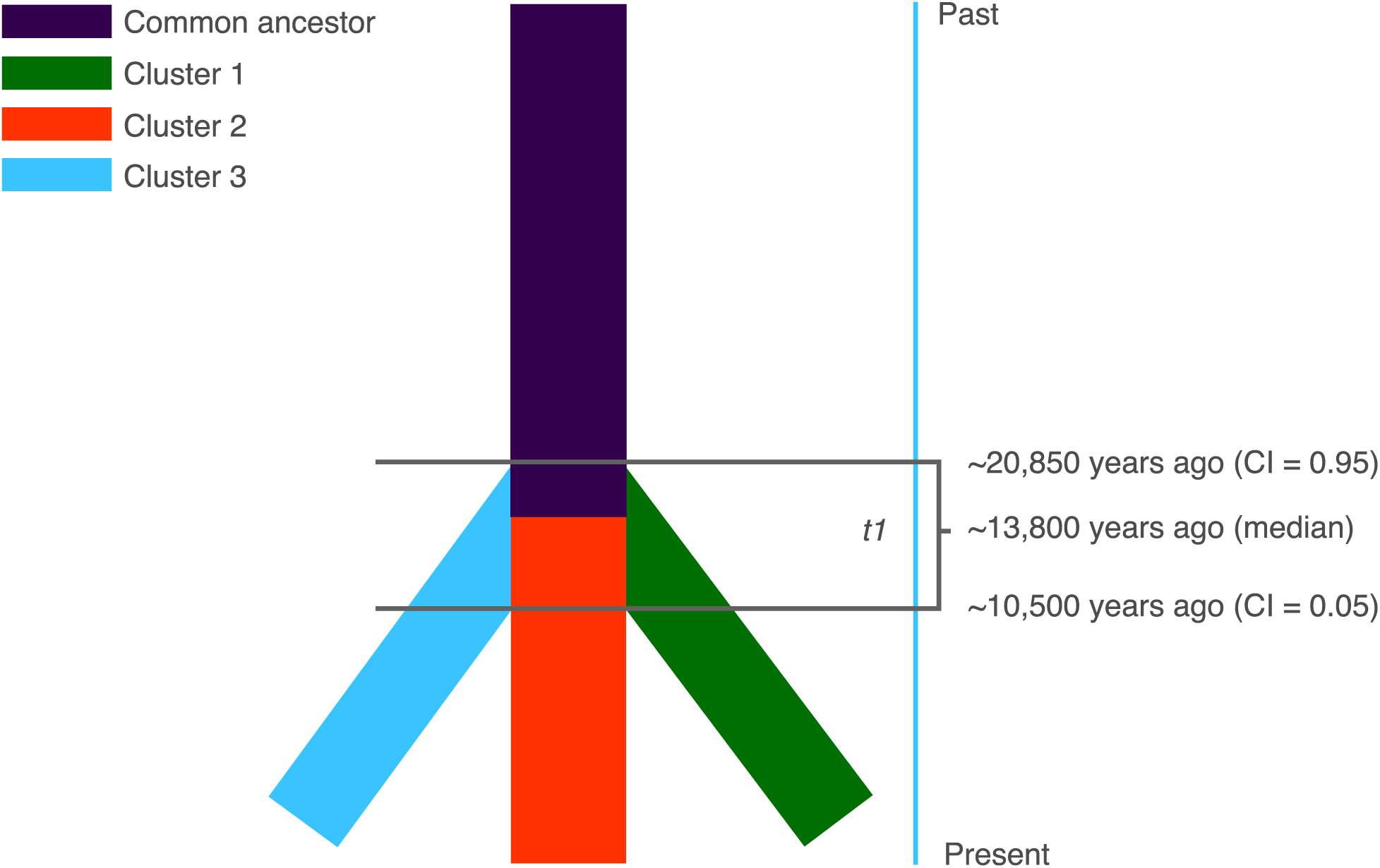
Most favored scenarios (Scenario 1) according to ABC-RF at the genepool level in the species’ core distribution. We defined three populations for this analysis: Cluster 1 includes Mbahambandara, Mabuni-buni, and Batuplat; Cluster 2 consists of Mamur, Kramongmongga, Pawada, Kokas, Nika, Ubadari, and Patimburak; and Cluster 3 is composed of Raduria, Perwasak and Purwahabtonggo. This scenario describes all three Cluster split at the same time during *t*1.

## DISCUSSION

### Influence of topographical barriers and past climate change on population dynamics and patterns of genetic diversity distribution

The genetic diversity within *M. argentea* is not homogeneously distributed throughout its range as demonstrated by all three types of genetic markers (nSSRs, cpDNA, and nrDNA). Moreover, our study also showed that certain populations within *M. argentea* native range have higher levels of genetic diversity than others.

Three main geographically-structured genetic clusters were observed within the native range of the species. The three genetic clusters corresponded to the following groupings of populations: South of Fakfak (pops 6), North East of Fakfak (pops 2 and 3), and North West of Fakfak (all other populations; Figure 2). This subdivision is further supported by the cpDNA data, and we hypothesized that these three genetic clusters might correspond to the joint action of topographic barriers and past climatic changes.

The region of Fakfak is crossed by the Fakfak Mountains (*Pegunungan Fakfak*) from East to West that represent a strong barrier to gene flow within the distribution of *M. argentea*. We observed a genetic differentiation between the North and South populations of the region, which probably support the role of Fakfak Mountains as a barrier to gene flow. Additionally, within the northern area of the Fakfak region, we have identified an East-West subdivision. The subdivision separates the populations of Batuplat and Mabuni-buni in the eastern part from the populations of Mamur, Kramongmongga, Pawada, Kokas, Nika, Ubadari, and Patimburak in the western part. There is a remarkable riverine network (the Kayuni River Network) in this region, which serves as a geographical midpoint between the two distinct genetic clusters. While the riverine barrier hypothesis has been extensively studied to assess genetic differentiation among populations (Gascon et al., 2000) and confirmed in the context of the Congo river network (Voelker et al., 2013), we did not anticipate the river to act as a barrier in this particular case. Actually, birds from the Bucerotidae family are the primary dispersers of Myristicaceae species (BBKSDA, 2022), and it is unlikely that river represent a constraint to their foraging activities. Consequently, we hypothesize that the Kayuni river network may have exerted a combined influence on gene flow patterns, alongside past climate fluctuations. This hypothesize was actually supported by our genetic structure and demographic analyses as discussed below.

The historical climate fluctuations were characterized by alternating periods of cooler climates, which significantly influenced the distribution and composition of forest refugia (De Frenne et al., 2021). During colder periods, the forest refugia likely contracted and became fragmented, isolating populations and promoting genetic differentiation (Connor, 1986). This situation might reflect the population structure in the two genetic clusters observed at the northern region of Fakfak Peninsula.

The influence of climatic events were also observed on the distinct evolutionary histories in the three genetic clusters within the species’ native range. The population inhabiting the northern coast of the Fakfak Peninsula exhibited a pattern of population expansion followed by a period of constant size. In contrast, the population in the southern coast of the peninsula displayed a trend of population decline. This demographic divergence coincides with the contrasting vegetation and topography between the north and south regions of the peninsula (Hill & Gleadow, 1989; Segar et al., 2017). The northern part is relatively flat and dominated by marshland, while the southern part is hilly and composed mainly of karst limestone (Balazs, 1968; Baldwin et al., 2012; Hill & Gleadow, 1989; Toussaint et al., 2021).

We found that the divergence of the three different clusters occurred approximately at the same time between ∼10,500 to ∼20,850 years ago. This timing of divergence is strikingly much more recent than the timing of demographic changes previously observed. The obtained divergence timings among clusters correspond to the climatic activities documented during the Late Pleistocene, specifically in Papua Island (Fairbairn et al., 2006). This period was characterized by the occurrence of multiple microclimatic conditions, as evidenced by sedimentation records (Lelono, 2008; Peng et al., 2021). These microclimatic shifts played a crucial role in shaping the vegetation formation in the mountainous range of Papua Island. Over time, the fluctuations in temperature and precipitation patterns influenced the distribution of plant species and impacted the composition of forest types found in the mountains. The dynamic nature of these climatic changes contributed to the rich biodiversity and distinct vegetation patterns observed in the region. Furthermore, the Fakfak Peninsula’s location, surrounded by seas on the north and south coasts with different monsoonal conditions (Prentice et al., 2007), may have contributed to the distinct evolutionary histories observed in the different populations of *M. argentea*.

Given the paucity of phylogeographic studies done in the region, we cannot compare our findings with other species. The development of a comparative phylogeographic framework would help investigating if similar patterns of genetic diversity distribution are found in other species, as well as additional studies on the demographic evolution of the species through time and of the timing of divergence of intra-specific clusters.

### History of introduction of the species

Our findings provide compelling evidence on the geographic origin of introduced populations in other regions of Papua Island (Manokwari and Raja Ampat) as well as the Moluccas (Ambon and Seram). The nuclear genetic analysis demonstrates that the introduced populations share the same genetic profile as some native populations, specifically Raduria, which is located in the southern coast of Fakfak.

Slightly different patterns were observed when examining the plastid compartment. In particular, the population in Manokwari did not share any haplotypes with the native range, whereas Ambon exhibited haplotype sharing with Batuplat and Raduria. Similarly, Seram displayed haplotypes shared with Raduria and Kokas. Overall, our results suggest that the Raduria population may have served as a main source of propagation material.

The observed pattern can be attributed to the migration of people from South Fakfak to various other regions of the island (Upton, 2009). The migration patterns discussed are strongly linked to the phenomenon of urbanization (Crosbie Walsh, 1987). The movement of tribal communities from rural areas in Fakfak to different regions of the island is primarily motivated by job prospects in Manokwari. The recent establishment of Manokwari as the capital of West Papua (Ronsumbre & Huwae, 2022), along with the growth of tourism in Raja Ampat (King, 2018), further contributed to this migration trend. It is plausible to speculate that during the migration process, individuals may have carried propagation materials with them to their new locations, enabling them to plant and utilize the species. Furthermore, it is worth noting that this human-mediated dispersal phenomenon extends beyond *M. argentea* and encompasses other forest species as well (Suryawan, 2016).

Finally, the intentional transportation of propagation materials during the human migration process indicates a deliberate effort to propagate and utilize various species in new locations. This finding emphasizes the significance of human activities in shaping the distribution of plant species within the region. Further research in this area may shed light on the specific mechanisms and impacts of human-mediated dispersal on local plant diversity.

## CONCLUSION

Our study revealed significant differences in genetic diversity between native and introduced populations of Papuan nutmeg. Specifically, several populations within the species’ native range exhibited higher genetic diversity compared to the introduced populations. The observed genetic clustering within the native range was influenced by geographical barriers and past climate changes, which likely contributed to the divergence of the three genetic clusters approximately 10,500 to 20,850 years ago. This divergence was likely facilitated by the presence of multiple microclimatic conditions. Considering these findings, it is imperative to implement sustainable management strategies for the genetic resources of *M. argentea*. One approach is to safeguard the distinct populations or gene pools by establishing specific conservation areas. The conservation efforts by local communities in the Fakfak Peninsula, serve as a valuable example that can be expanded and applied on a larger scale by policymakers. It is important to emphasize that these conservation efforts also contribute to the maintenance of genetic resources for these forest species. By effectively communicating the benefits of conserving genetic diversity, we can further promote and enhance these conservation initiatives.

## ACKNOWLEDGEMENT

The authors express their gratitude to the farmers and landowners of Ambon, Seram, Raja Ampat, Manokwari, and Fakfak for generously providing valuable information on the history of Papuan nutmeg and for their assistance in collecting leaf samples. Additionally, we extend our appreciation to our guide and Dina Tri Wahyuni for their valuable support during the field work. We also thank Roxali Bijmoer and Marnel Scherrenberg from Naturalis Biodiversity Center, Leiden, The Netherlands as well as Myriam Gaudeul from National Museum of Natural History, Paris, France for their invaluable help during the specimen collection. We further thank Pierre Mournet from CIRAD, Montpellier, France and Sylvain Santoni, Audrey Weber from INRAe, Montpellier, France for providing assistance during lab analysis. This study was authorized by Indonesian National Research and Innovation Agency (BRIN) with research permit to JD (144/SIP/FRP/E5/Ditk.KI/V/2018) for collecting samples in Ambon, and collecting permit to JK (B-3382/IPH.1/KS.02.04/IX/2019 and B-75/IV/KS.01.04/3/2022). We also greatly appreciate the support from the Indonesia Endowment Fund for Education (LPDP) to JK with grant No. 0003719/AFR/D/BUDI-2018. This project also supported by Agropolis Fondation under the reference ID 1502-503 and 1502-504 through the « *Investissements d’avenir* » program (Labex Agro: ANR-10-LABX-0001-01), under the frame of I-SITE MUSE (ANR-16-IDEX-0006). The authors acknowledge the ISO 9001 certified IRD i-Trop HPC (member of the South Green Platform) at IRD Montpellier for providing HPC resources that have contributed to the research results reported within this paper. URL: https://bioinfo.ird.fr/- http://www.southgreen.fr.

## Supplementary information

**Table S1.** Description of the historical parameter priors on the demographic inference of *M. fragrans* in ABC analysis.

| Parameter type | Parameter name | Prior distributions |
| --- | --- | --- |
| Population size<sup>†</sup> | Ne | Log_U [10 - 50,000,000) |
|  | N1 | Log_U [10 - 1,000,000] |
|  | N2 | Log_U [10 - 25,000,000] |
|  | N3 | Log_U [10 - 5,000,000] |
| Divergence time<sup>‡</sup> | $t1$ | U [10 - 1000] |
| | $t2$ | U [10 - 10000] |
| Conditions | $t2 > t1$ | |

<sup>†</sup>In number of reproductively effective diploid individuals. <sup>‡</sup>In number of generations, assuming a generation time of × years. Distributions were considered either uniform (U) or Log-uniform (Log\_U). Reference tables were constructed for 100,000 data sets per scenario.

**Figure S1.**
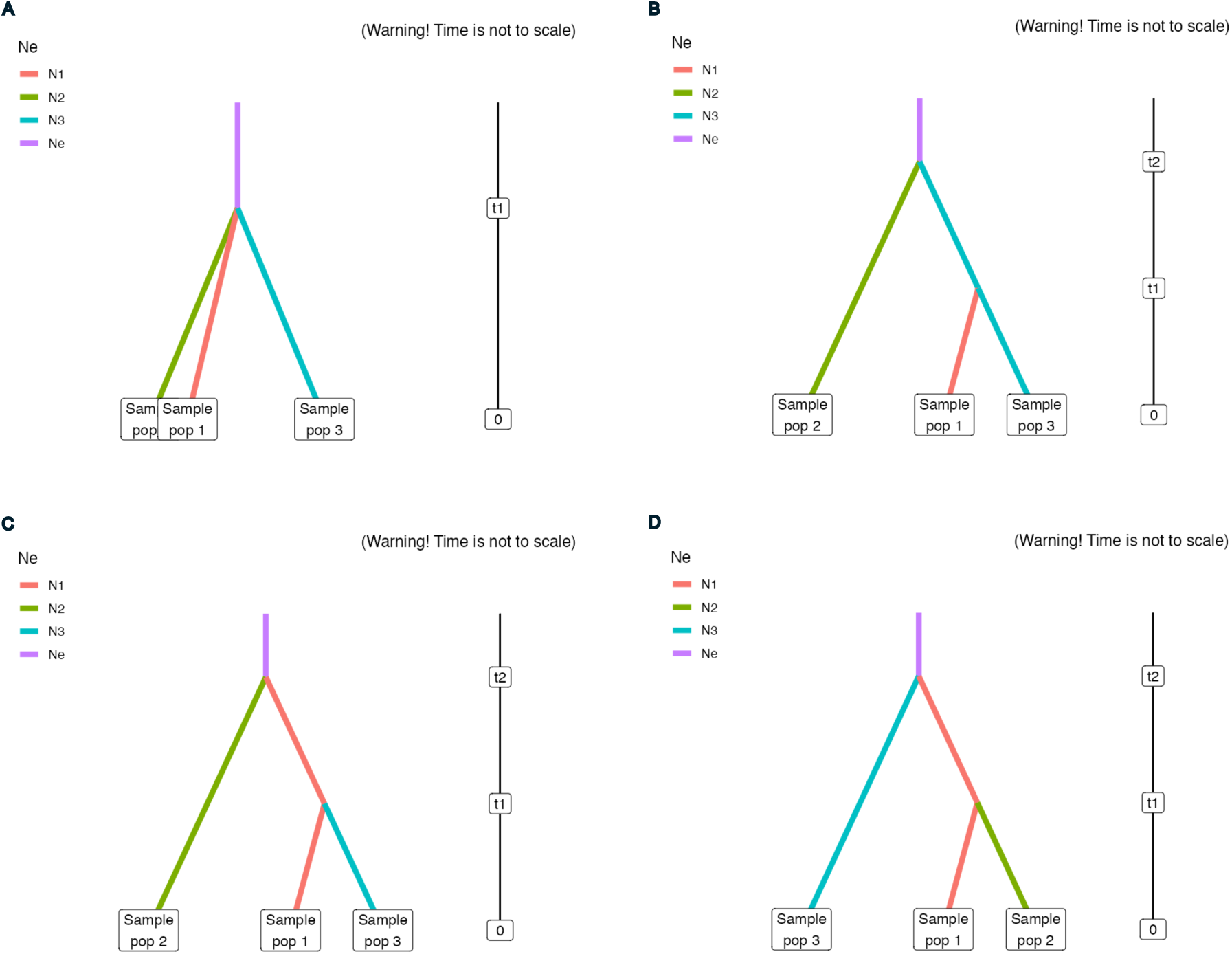
Graphic representation of inferred population history of *M. argentea*. The three distinct genepool of *M. argentea*’s core distribution was based on STRUCTURE analysis. The aim is to identify the ancestral population genetic cluster that serveed as the basis for the genetic clusters outside of its native range. We defined three populations for this analysis: Pop 1 includes Mbahambandara, Mabuni-buni, and Batuplat; Pop 2 consists of Mamur, Kramongmongga, Pawada, Kokas, Nika, Ubadari, and Patimburak; and Pop 3 is composed of Raduria, Perwasak and Purwahabtonggo. We proposed four scenarios to infer demographic history using ABC analysis. In Scenario 1, all three Pops split at the same time at *t*1. Scenario 2 suggests that Pop 2 and Pop 3 diverged at *t*2 and Pop 1 originated from Pop 3 at *t*1. In Scenario 3, Pop 1 and Pop 2 diverged at *t*2 and originated from Pop 1 at *t*1. Finally, in Scenario 4, Pop 1 and Pop 3 diverged at *t*2 and Pop 2 originated from Pop 1 at *t*1.

**Figure S2.**
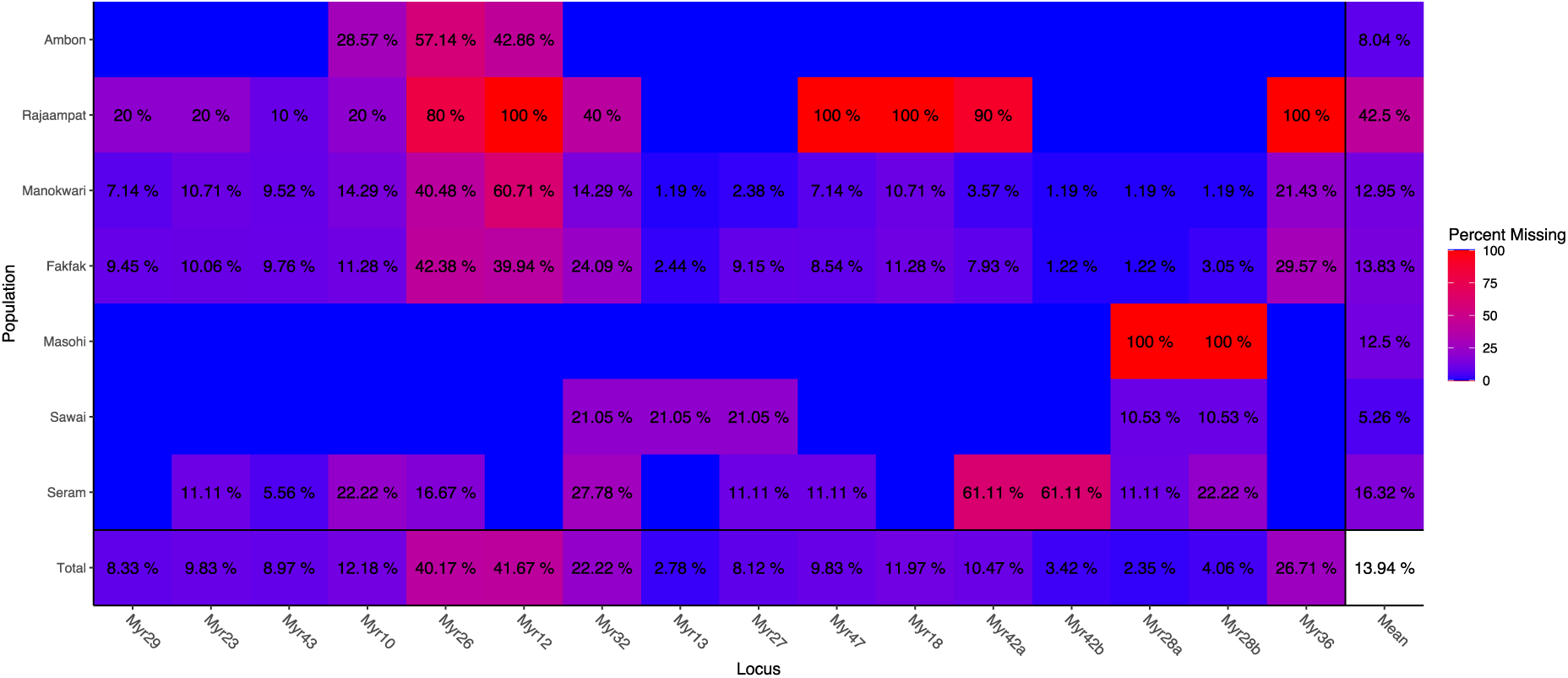
Percentage of missing data of 473 *M. argentea* individuals in 5 different populations as inferred using 16 nSSRs. Masohi and Sawai are located in a same island as Seram population, we therefore grouped them together.

**Figure S3.**
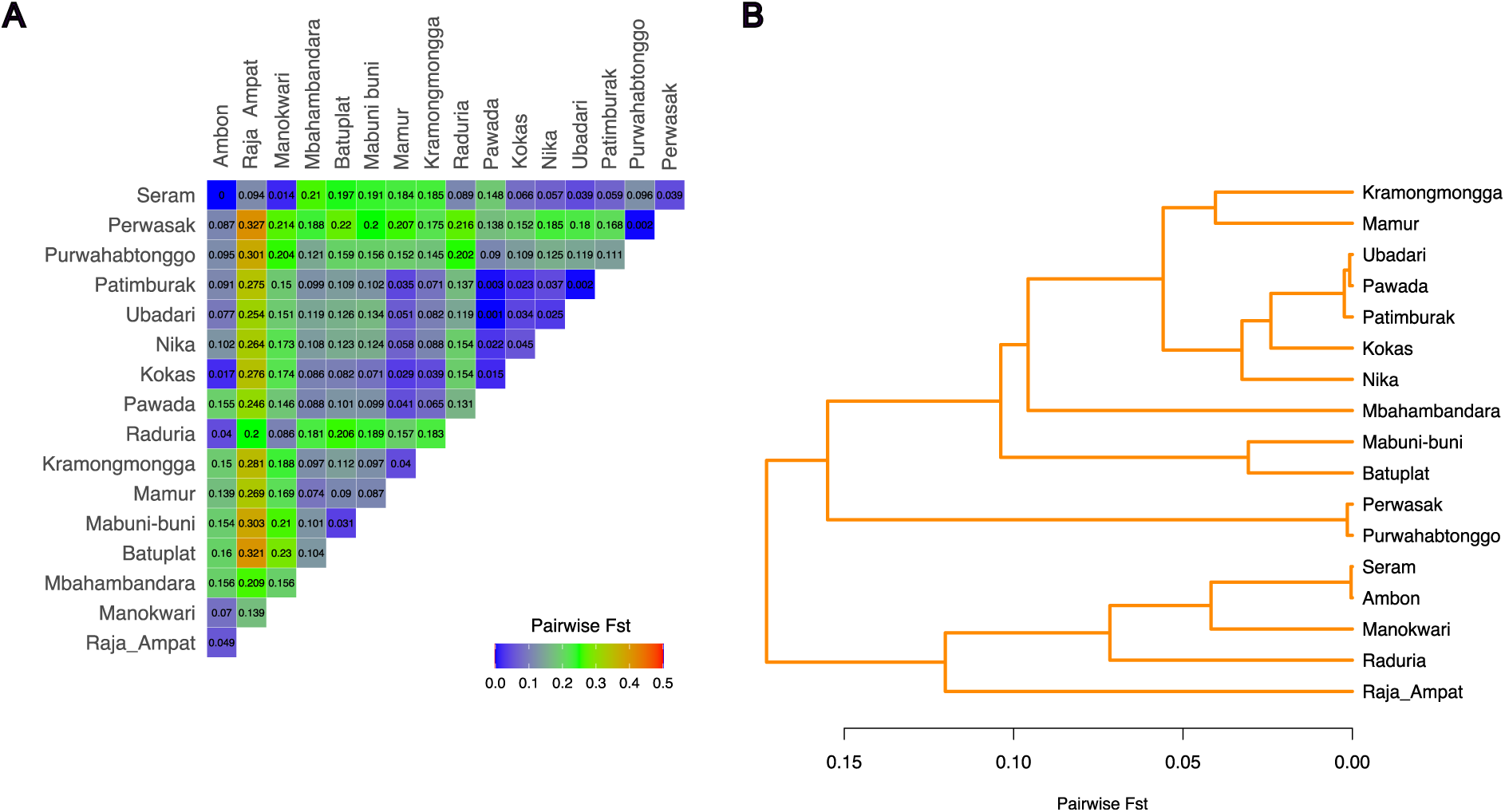
Pairwise *F*_ST_ in all populations inferred by SSR. Values are visualized in a heatmap plot (A) and a dendrogram tree (B) in all thirteen populations.

**Figure S4.**
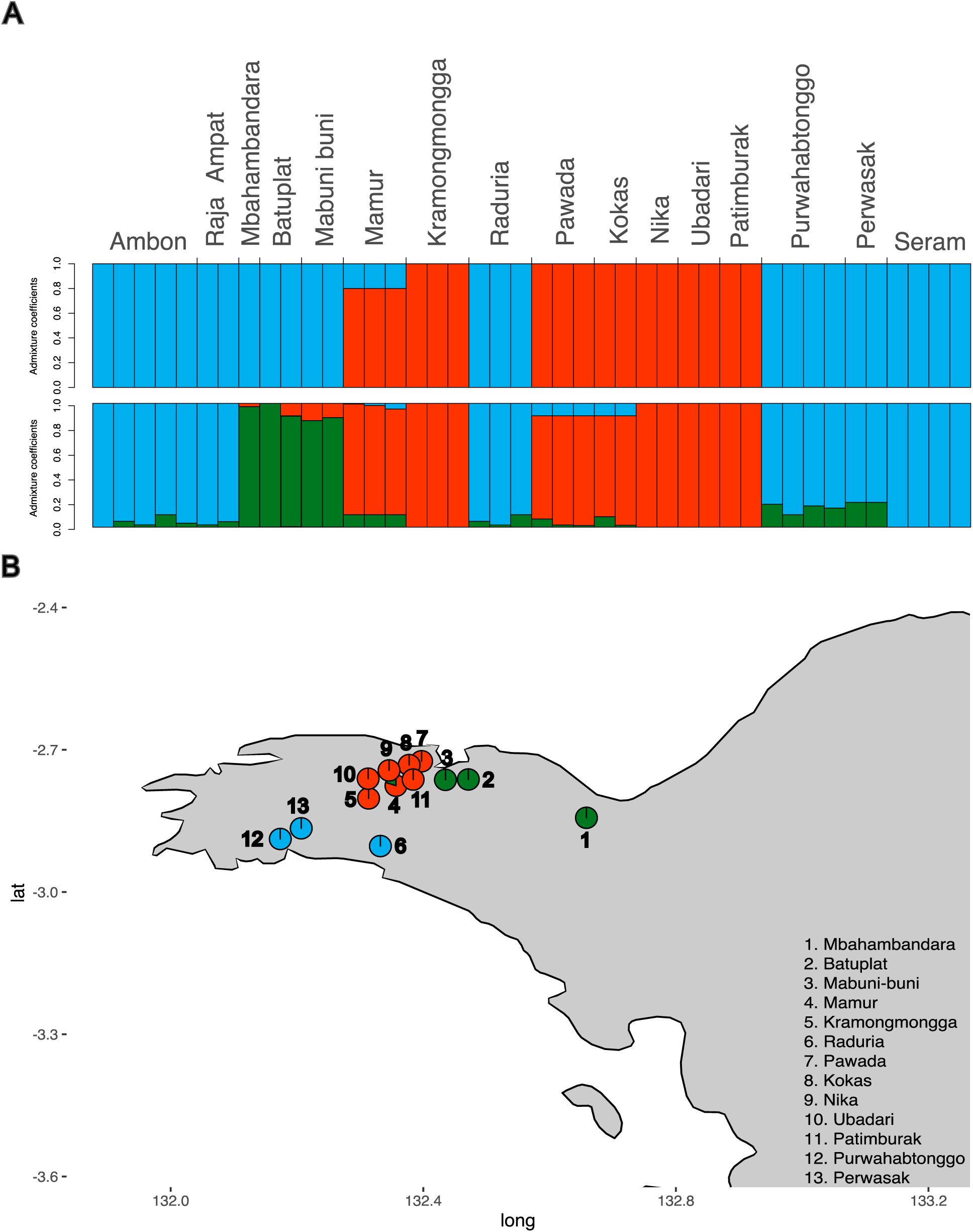
Population structure in all populations using fastSTRUCTURE. Barplots of 256 individual admixture proportions, for *K* = 2 and *K* = 3 (A). Geographic projection of *K* = 2 in 257 the species’ core distribution, Fakfak Peninsula, Papua Island, Indonesia (B).

**Figure S5.**
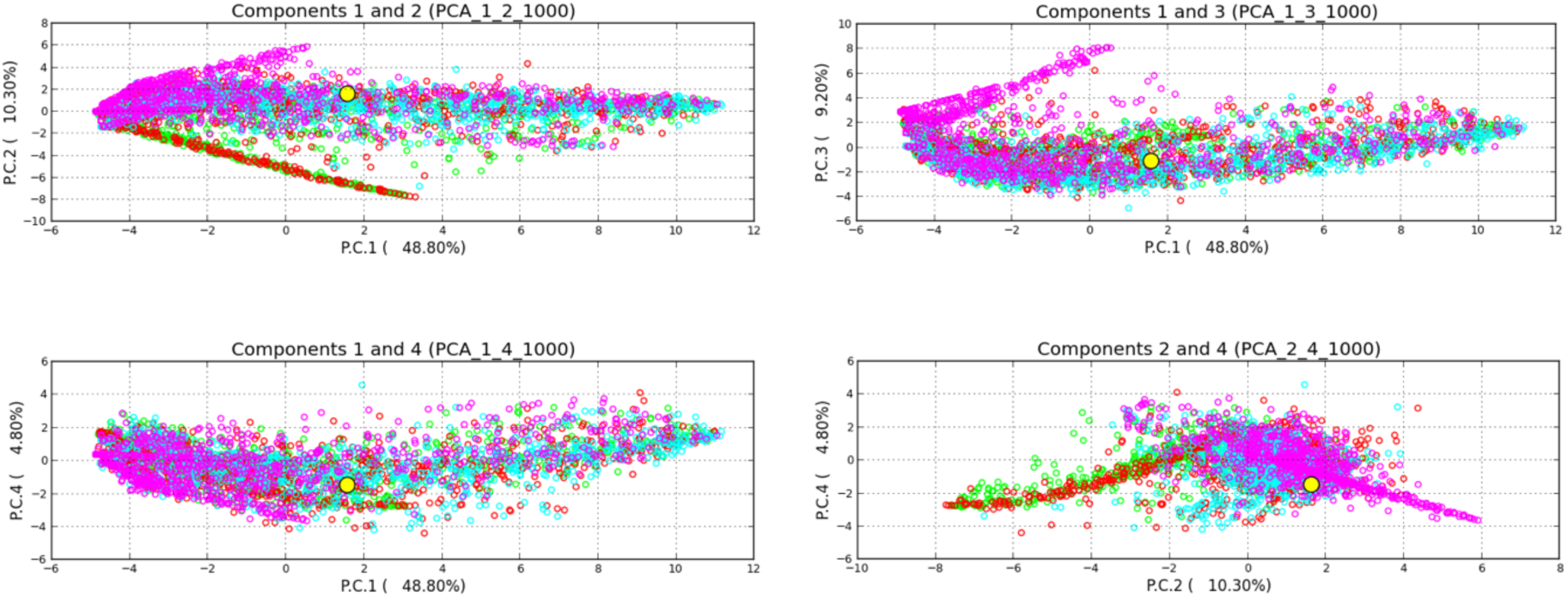
Principal Component Analysis (PCA) of the prior-scenario combinations and the observed dataset using DNA sequence. Scenario 1, 2, 3, and 4 priors are indicated with green, blue, purple, and red dots respectively, and observed dataset is indicated by a yellow dot.

